# The CXCR6/CXCL16 axis links inflamm-aging to disease severity in COVID-19 patients

**DOI:** 10.1101/2021.01.25.428125

**Authors:** Daniel J. Payne, Surita Dalal, Richard Leach, Richard Parker, Stephen Griffin, Clive S. McKimmie, Graham P. Cook, Stephen J. Richards, Peter Hillmen, Talha Munir, Louise Arnold, Kathryn Riley, Claire McKinley, Sandra Place, Richard L. Baretto, Darren J. Newton

**Author notes:** Corresponding author: Mr D Payne, Consultant Clinical Scientist, University Hospitals of Leeds, St James Hospital, Leeds UK. The authors have declared that no conflict of interest exists.

## Abstract

Advancing age and chronic health conditions, significant risk factors for severe COVID-19, are associated with a pro-inflammatory state, termed inflamm-aging. CXCR6^+^ T cells are known to traffic to the lung and have been reported to increase with age. The ligand of CXCR6, CXCL16, is constitutively expressed in the lung and upregulated during inflammatory responses and the CXCR6/CXCL16 axis is associated with severe lung disease and pneumonia. Genome-wide association studies have also recently identified 3p21.31, encompassing the *CXCR6* gene, as a susceptibility locus for severe COVID-19. We assessed numbers T cells expressing the chemokine receptor CXCR6 and plasma levels of CXCL16, in control and COVID-19 patients. Results demonstrated that circulating CD8^+^CXCR6^+^ T cells were significantly elevated with advancing age, yet virtually absent in patients with severe COVID-19. Peripheral levels of CXCL16 were significantly upregulated in severe COVID-19 patients compared to either mild COVID-19 patients or SARS-CoV-2 negative controls. This study supports a significant role of the CXCR6/CXCL16 axis in the immunopathogenesis of severe COVID-19.

## Introduction

Coronavirus disease 2019 (COVID-19), caused by infection with severe acute respiratory syndrome coronavirus-type 2 (SARS-CoV-2), encompasses clinical phenotypes ranging from asymptomatic infection, through to severe disease and death. The more severe end of this spectrum is often associated with respiratory pathology (1, 2). It is well established that the course of any infection is dependent on a number of variables including pathogen virulence, environmental and host factors. The latter includes variation in the immune response driven by genetics, age and the presence of co-morbidities. SARS-CoV-2 infection induces both innate and adaptive immunity (3) and severe COVID-19 is associated with exaggerated T cell responses producing increased levels of pro-inflammatory cytokines including IL-6, TNF-α, and IL-1 (4, 5). This aggressive hyper-inflammatory state results in significant lung damage and high mortality (4). Post-mortem studies have shown both lymphocyte and neutrophil lung infiltration, indicating that migration of pro-inflammatory cells into the lung is a key step in the pathology and outcome of this infection (6, 7). Immunomodulatory therapies, including the anti-IL-6 monoclonal antibody tocilizumab and the corticosteroid dexamethasone, may improve outcomes, highlighting the importance of inflammatory processes in COVID-19 pathology (8, 9).

Current epidemiological studies have identified advancing age, chronic health conditions, such as diabetes and obesity, and certain ethnicities as risk factors for more severe disease (10). Advancing age has been strongly associated with a pro-inflammatory immune phenotype, so-called inflamm-aging, where T cells acquire a more innate NK cell-like pro-inflammatory phenotype associated with upregulation of markers of both T cell exhaustion and senescence (11). Furthermore, increased expression of chemokine receptors, including CXCR6 on T cells, has been demonstrated in aging animal models (12). The receptor CXCR6 (CD186) is expressed on activated T cells, NK cells, NKT cells and mucosal-associated invariant T (MAIT) cells (13–15).

## Results and Discussion

We assessed the expression of CXCR6^+^ on CD4^+^ and CD8^+^ T cells in consecutive blood samples to determine age-related differences and thereby support a potential role of these cells in inflamm-aging and justify further assessment in the pathogenesis of COVID-19 (16–18) (Figure 1).

**Figure 1.**
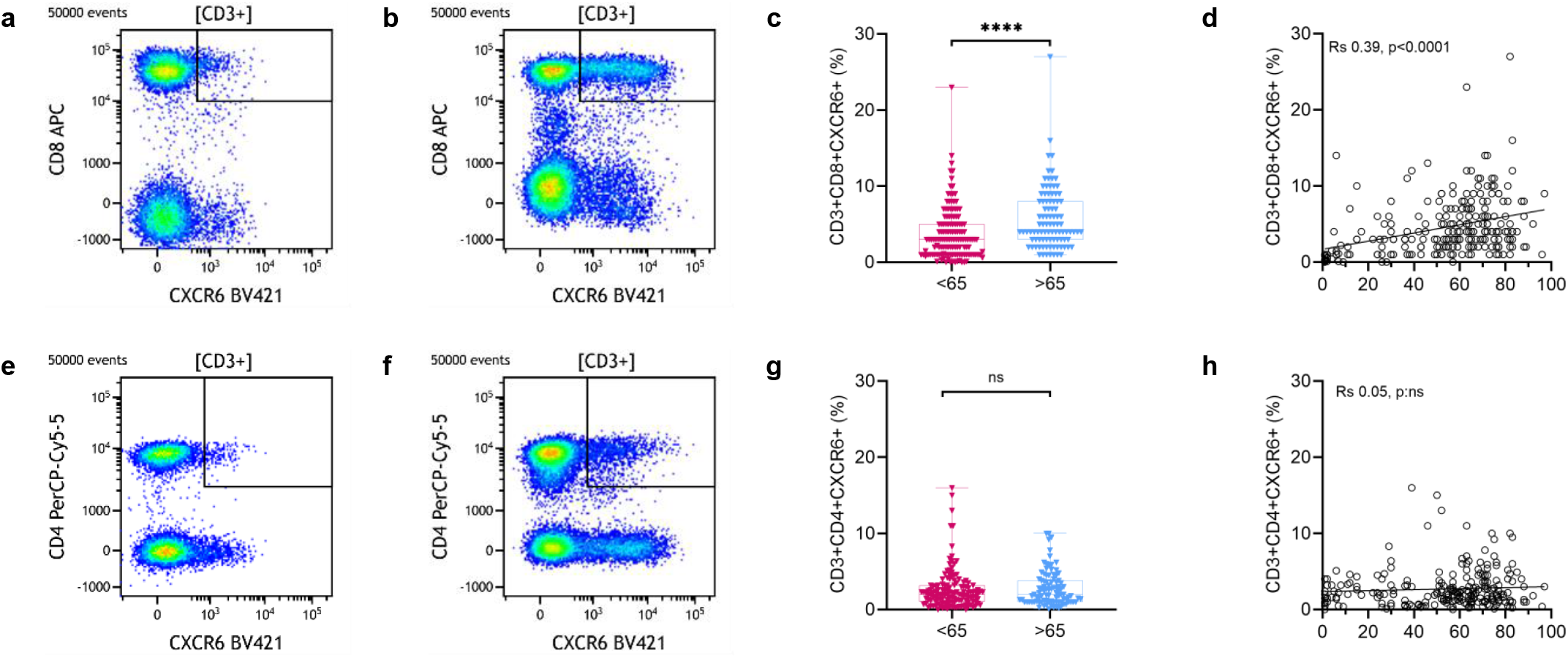
Age-related percentage of CXCR6^+^ T cells in consecutive samples. SARS-CoV-2 status of these samples was undetermined. Data presented as a percentage of CD3+ gated T cells. Dot plots illustrate differences observed in proportion of CD3^+^CD8^+^CXCR6^+^ cells **a:** 15 year old **b:** 83 year old. **c:** Levels of CD3^+^CD8^+^CXCR6^+^ cells in peripheral blood of different age groups **(<65)** <65 years old (n=137). **(>65)** >65 years old (n=96). Median, maximum, and minimum values shown. Mann Whitney was used to compare populations, **** p<0.0001;. **d:** Correlation of CD3^+^CD8^+^CXCR6^+^ with age in 233 peripheral blood samples. Spearman rank correlation: R_s_ 0.39, p<0.0001, suggesting a trend to increase with age. Dot plots illustrate differences observed in proportion of CD3^+^CD4^+^CXCR6^+^ cells **e:** 15 year old and **f:** 83 year old. **g:** Levels of CD3^+^CD4^+^CXCR6^+^ cells in peripheral blood of different age groups **(<65)** <65 years old (n=137). **(>65)** >65 years old (n=96). Median, maximum, and minimum values shown. Mann Whitney was used to compare populations, ns: not significant. **h:** Correlation of CD3^+^CD4^+^CXCR6^+^ with age in 233 peripheral blood samples: R_s_ 0.05, p: ns.

Patients over 65 years of age are known to have more severe outcomes in COVID-19 (19). In keeping with the inflamm-aging hypothesis, we demonstrated a highly significant increase in CD8^+^CXCR6^+^ T cells in the blood of patients aged over 65 years (n=96) compared to those aged under 65 years (n=137) (p<0.0001; Figure 1c). A progressive increase with advancing age was also observed (Rs 0.39, p<0.0001; Figure 1d). There were lower proportions of CD4^+^CXCR6^+^ T cells were observed compared to CD8^+^CXCR6^+^ T cells and there were no significant age-related differences (Figure 1g and 1h). The increased frequency of CD8^+^CXCR6^+^ T cells in the blood of older patients is supportive of a pro-inflammatory phenotype, potentially rendering this group susceptible to hyper-inflammatory immune responses associated with poor outcomes in COVID-19.

Peripheral blood T lymphopenia has been identified as an immunological marker for SARS-CoV-1 and 2 infection with postulated mechanisms including immune-mediated destruction and trafficking to pathological sites (20–22). We analysed the absolute numbers and proportion of CD4^+^ T cells, CD8^+^ T cells and NK cells relative to total leucocytes in controls, mild and severe COVID-19 patients. There was an absolute and proportional reduction in CD4^+^ and CD8^+^ T cells in severe COVID-19 (Figure 2). This was more pronounced for CD8^+^ T cells, with statistical significance in severe COVID-19 compared to controls (p<0.001 and p<0.0001 for absolute number and proportion respectively). There was no significant difference in NK cells.

**Figure 2.**
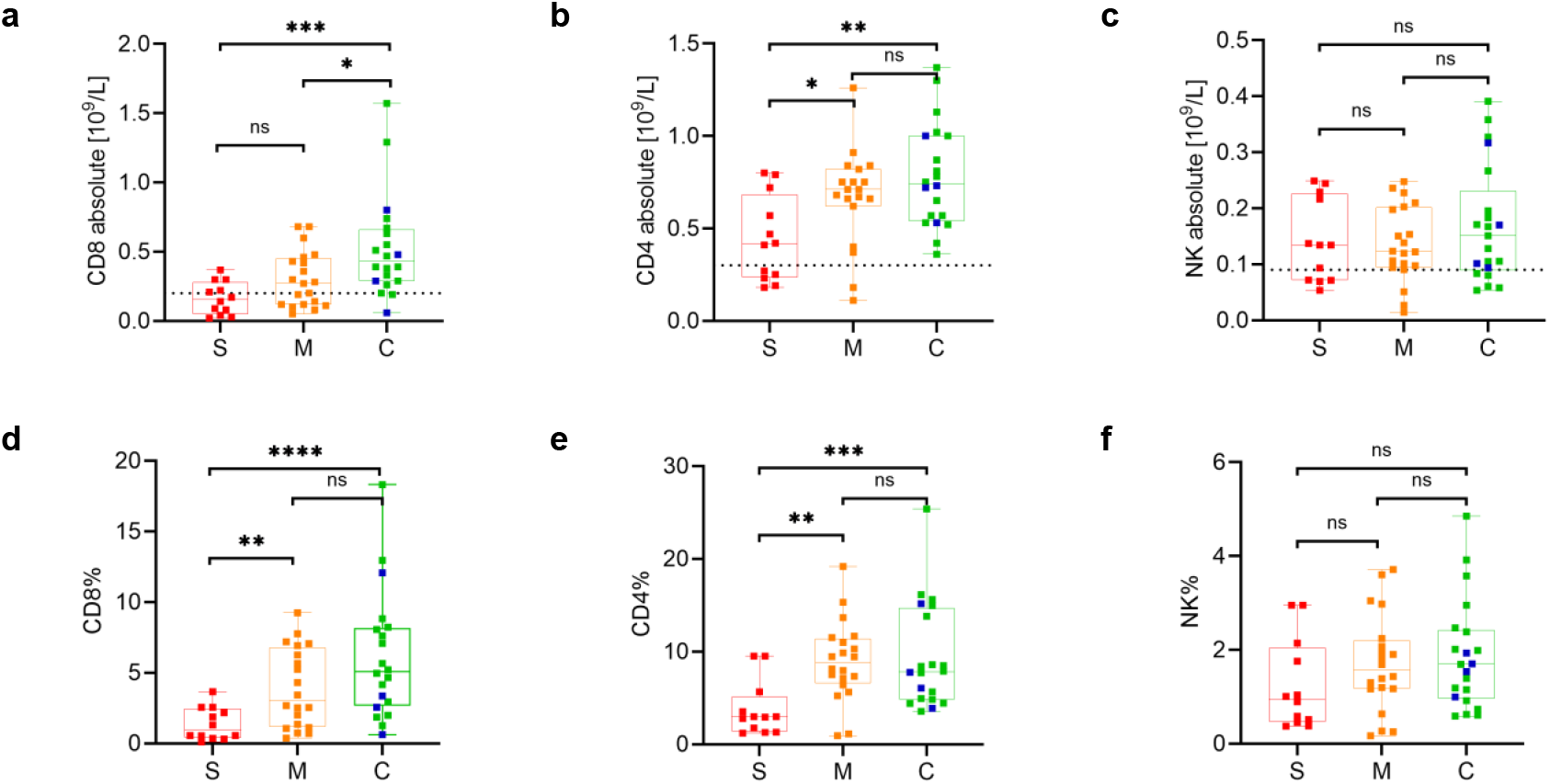
Reduced absolute and total T-cell numbers in severe COVID-19 patients. **(S)**<u>evere COVID-19</u>; SARS-CoV-2 RT-PCR positive patients on ITU (n=12, red). **(M)**<u>ild COVID-19</u>; SARS-CoV-2 RT-PCR positive patients non-ITU (n=20, orange). **(C)**<u>ontrols</u>; SARS-CoV-2 RT-PCR negative non-ITU (n=16, green). <u>on ITU</u> (n=4, blue). maximum, and minimum values shown, dotted line shows lower end of absolute reference range. Mann Whitney was used to compare populations; **** p<0.0001 *** p<0.001 ** p<0.01 * p<0.1 ns = not significant. **a** and **d:** Significantly reduced absolute and percentage CD8^+^ T cells in when comparing S to M or C. **b** and **e:** Significantly reduced absolute and percentage CD4^+^ T cells in when comparing S to M or C. **c** and **f:** No significant difference in NK cells was observed when comparing S, M to C.

The receptor for SARS-CoV-2 is angiotensin converting enzyme 2 (ACE2) which is highly expressed on alveolar epithelial type II cells of the lower respiratory tract (23). Membrane bound CXCL16 is constitutively expressed on bronchial epithelial cells and is released in metalloprotease-dependent manner in an inflammatory environment, producing a soluble form which is chemotactic for CXCR6^+^ T-cells (24, 25). The CXCR6/CXCL16 axis mediates homing of T cells to the lungs in disease (26–28) and hyper-expression is associated with localised cellular injury (29–31). Murine studies have demonstrated that this axis is involved in lung pathology associated with other infections, including influenza, with antagonism resulting in reduced tissue inflammation (26, 29).

CXCL16 is up-regulated during viral infections and mediates CD8^+^CXCR6^+^ T-cell recruitment (32, 33). Differential expression of *CXCR6* and *CXCL16* mRNA was observed in severe COVID-19 compared to mild disease (34) and significant functional polymorphisms in *CXCR6* are linked to viral control (35). Furthermore, in HIV infection *CXCR6* polymorphisms have been linked with certain ethnicities associated with more severe lung pathology and poorer outcomes (36). We compared peripheral blood T cell populations in severe and mild COVID-19 to control samples. This revealed that absolute CD8^+^ CXCR6^+^ T cell populations were significantly reduced in both severe and mild COVID-19 patients compared to controls (p<0.0001 and p<0.1 respectively; Figure 3e), with significant reduction in absolute CD4^+^ CXCR6^+^ T cells only between severe COVID-19 and controls (p<0.001; Figure 3g). Strikingly, both CD4^+^ and CD8^+^ CXCR6^+^ expressing T cells were present at extremely low proportions in the blood of severe COVID-19 patients (n=12).

**Figure 3.**
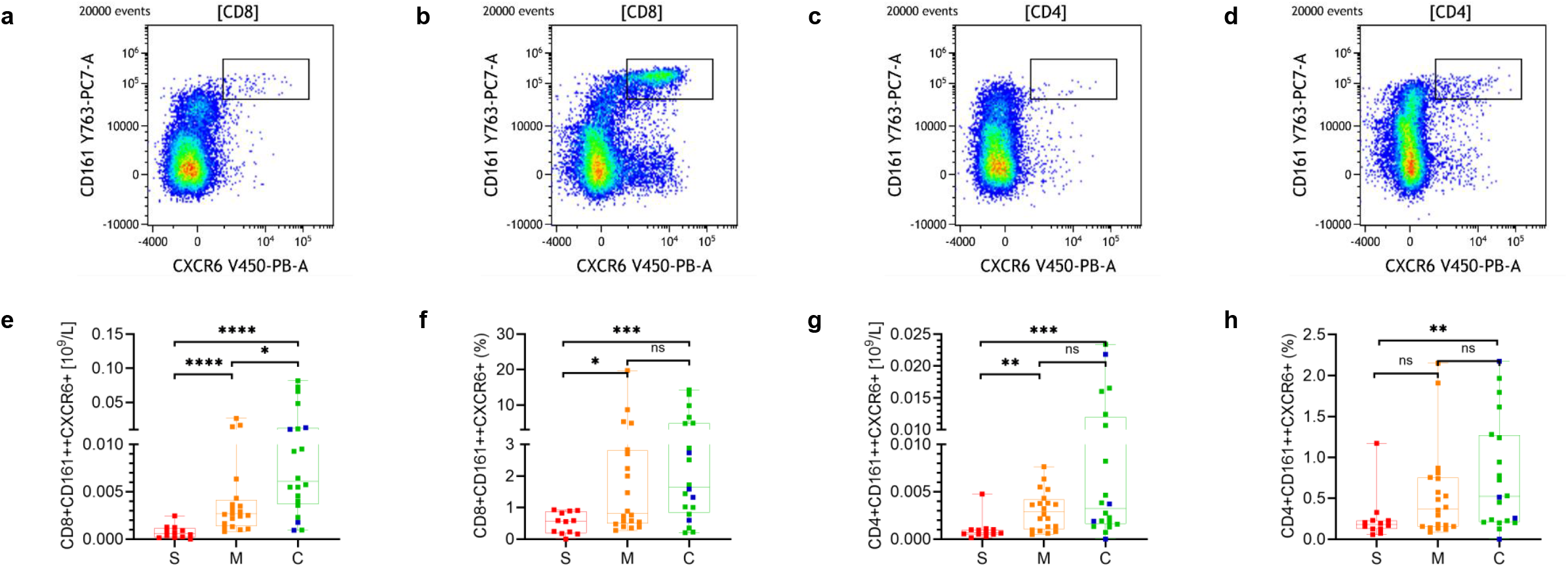
Reduced CXCR6^+^T cells in severe COVID-19 patients. Gated on CD3^+^CD8^+^ T cells, **a:** illustrates reduced CXCR6^+^CD161^++^ cells in severe COVID-19 when compared with **b:** control sample. Gated on CD3^+^CD4^+^ T cells, **c:** illustrates reduced CXCR6^+^CD161^++^ cells in severe COVID-19 when compared with **d:** control sample. **e:** Significantly reduced absolute and **f:** percentage of CD8^+^CD161^++^CXCR6^+^ cells in severe COVID-19 compared to mild COVID-19 and controls. **e:** Significantly reduced absolute and **f:** percentage of CD4^+^CD161^++^CXCR6^+^ cells in severe COVID-19 compared to mild COVID-19 and controls. **(S)**<u>evere COVID-19</u>; SARS-CoV-2 RT-PCR positive patients on ITU (n=12, red). **(M)**<u>ild COVID-19</u>; SARS-CoV-2 RT-PCR positive patients non-ITU (n=20, orange). **(C)**<u>ontrols</u>; SARS-CoV-2 RT-PCR negative non-ITU (n = 16, green), <u>on ITU</u> (n=4, blue). Median, maximum, and minimum values shown. Mann Whitney was used to compare graphed populations. **** p<0.0001 *** p<0.001 ** p<0.01 * p<0.1 ns = not significant

Studies in COVID-19 patients suggest CXCR6 correlates directly with the proportion of MAIT cells and that this population is reduced in the peripheral blood in COVID-19 (37). Killer cell lectin like receptor subfamily B, member 1 (CD161) is a C-type lectin receptor expressed on NK cells and a subset of T cells with both stimulatory and inhibitory functions. Expression of this marker was assessed on CD8^+^ T cells as high levels of expression have been associated with MAIT cells and Th17 responses (38). We demonstrated that the majority of CD3^+^ CD8^+^ CXCR6^+^ T cells in controls and mild COVID-19 cases were CD161^++^ CD45RA^-^ CD27^+^ HLA-DR^-^ CD57^-^ (Figure 3 and Supplementary Figure 1) suggestive of an effector memory profile. In most cases these cells were also positive for CD56 and CD279 (see Supplementary Figure 1), consistent with either NKT cells, invariant T cells or mucosal-associated invariant T (MAIT) cells (13). CD8^+^ CXCR6^+^ CD161^-^ T cells were present in lower numbers and there was no significant difference in this population between COVID-19 patients and controls. Proportions of both CD4^+^ CXCR6^+^ CD161^++^ and CD4^+^ CXCR6^+^ CD161^-^ T cells were lower compared to CD8^+^ T cells. The extended phenotype of the CD4^+^ CXCR6^+^ CD161^++^ is suggestive of an effector/central memory population (Supplementary Figure 1).

Cells with this phenotype have been shown to exhibit tissue homing properties, with infiltrates described in inflammatory diseases including rheumatoid arthritis, psoriasis, multiple sclerosis and Crohn’s disease (39). Levels of circulating and pancreatic MAIT-cells in type 1 diabetes mirror the findings of our MAIT-like cells in COVID-19. In type 1 diabetes high levels of circulating cells are present at diagnosis with numbers falling after 1 year with a concurrent increase in pancreatic numbers suggesting trafficking and a potential pathogenic role (40).

To further characterise the CXCR6/CXCL16 axis in the immunopathogenesis of COVID-19, plasma concentrations of CXCL16 from 28 COVID-19 patients and 12 controls were assessed. CXCL16 was significantly elevated in severe COVID-19 samples (n=10) when compared with mild COVID-19 (n=18, p<0.0001) or controls (n=12, p <0.001; Figure 4a), which contrasts with the findings of Liao *et al* (41) where *CXCL16* mRNA was more highly expressed in bronchoalveolar lavage fluid in mild disease, albeit with significantly fewer patient numbers in all groups. There was an inverse relationship between the concentration of blood CXCL16 and the proportion of CD8^+^ and CD4^+^ CXCR6^+^ T cells in the blood in COVID-19 patients (Figures 4b and 4c). This suggests trafficking of CXCR6^+^ T cells to the lung drives a pro-inflammatory immunopathology in severe COVID-19, with these cells infiltrating into the tissue, which is supported by lower numbers of CD8^+^ T cells reported in broncho-alveolar lavage in mild compared to severe COVID-19 patients (41). Furthermore, the CXCR6/CXCL16 axis has been implicated in both infective (influenza) and non-infective (sarcoidosis) inflammatory lung diseases (25, 26). However, this inverse relationship between CXCL16 levels and CXCR6^+^ T cells and may also be explained by either CXCL16 binding to CXCR6 causing receptor internalisation, epitope masking or CXCL16-mediated T cell apoptosis.

**Figure 4.**
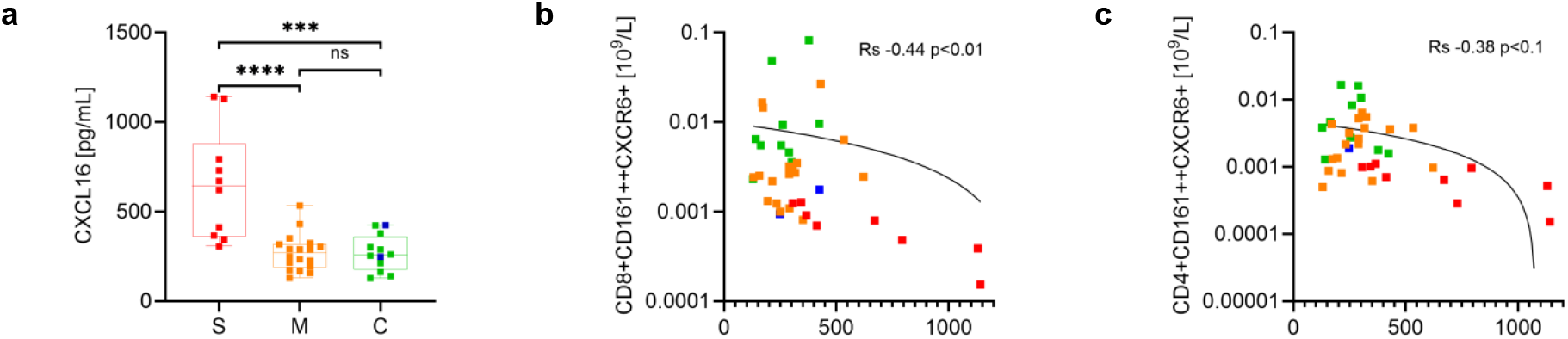
Plasma concentrations of CXCL16 in COVID-19 compared with controls. **a:** Median, maximum, and minimum values shown. **(S)**<u>evere COVID-19</u>; SARS-CoV-2 RT-PCR positive patients on ITU (n=10, red). **(M)**<u>ild COVID-19</u>; SARS-CoV-2 RT-PCR positive patients non-ITU (n=18, orange). **(C)**<u>ontrols</u>; SARS-CoV-2 RT-PCR negative non-ITU (n=10, green). <u>on ITU</u> (n=2, blue). (**** p<0.0001 *** p<0.001 ns = not significant) Significantly increased levels of CXCL16 are present in the plasma of severe COVID-19 patients. **b** and **c:** CD8^+^ and CD4^+^ CD161^++^CXCR6^+^ T cell count falls (log values) as plasma concentration of CXCL16 increases, SARS-CoV-2 RT-PCR positive patients on ITU (n=10, red), SARS-CoV-2 RT-PCR positive patients non-ITU (n=18, orange), SARS-CoV-2 RT-PCR negative non-ITU (n=10, green). <u>on ITU</u> (n=2, blue), line logistical regression shown. Spearman rank correlation: R_s_ −0.44, p<0.01 and R_s_ −0.38, p<0.1 for CD8^+^ and CD4^+^ cells respectively

Following infection with SARS-CoV-2, there is potential for pre-existing inflammatory CXCR6^+^ populations, associated with either co-morbidity and/or inflamm-aging, to be recruited from the blood to the lungs mediated by CXCL16, resulting in more severe disease (42). Similarly, in other diseases characterised by a T cell infiltrate, such as type 1 diabetes, high expression of this chemokine receptor and ligand have been reported in pancreatic tissue where they play a role in inflammation (43).

CD8^+^CD161^++^CXCR6^+^ T cells, have the capacity to be cytotoxic and express the transcription factor RORγt, which is associated with a Th17-like phenotype and a pro-inflammatory cytokine profile (IFN-γ, TNF-α, IL-17 and IL-22) along with expression of cytotoxic mediators such as granzyme (14, 44). These factors have all been shown to be significantly elevated in severe, but not mild COVID-19 patients, despite a more profound lymphopenia (45, 46). These cells have also been implicated in other lung infections, with IL-17 mediated inflammation and pathogenesis reported in patients with immune-mediated community-acquired pneumonia (47). A similar Th17 profile has been described in patients with COVID-19(48) along with a significant reduction in circulating CD161^++^ cells (48, 49). It is likely that these CD161^++^ cells are identical to the T cells identified in this study. As well as mediating chemotaxis of inflammatory cells, murine studies suggest that elevated levels of CXCL16 may directly contribute to lung injury through production of reactive oxygen species and compromised epithelial barrier integrity, with CXCL16 inhibitors protecting against lipopolysaccharide-mediated lung injury (29).

This study demonstrates an age-related increase in CD8^+^CXCR6^+^ T cells consistent with inflamm-aging in humans and that more severe outcomes in COVID-19 associate with increased peripheral CXCL16 and reduced circulating CXCR6^+^ T cells, suggesting an immunopathogenic role. This may have significant implications in the stratification of the risk for patients infected with SARS-CoV-2 and raises the possibility of novel therapeutic agents targeting this axis in severe COVID-19. Studies on CXCR6 expression on T-cells and levels of CXCL16 in dexamethasone and tocilizumab treated patients will provide further insight into the pathogenesis and putative mechanisms of therapy. This axis may also be relevant in other infections associated with lung pathology such as influenza.

## Methods

### Samples

#### Consecutive samples

CXCR6 expression was assessed on CD4+ and CD8+ T cells as part of routine diagnostics on samples taken between 30th March and 1st July 2020 from patients not tested for SARS-CoV-2. Samples consisted of 233 peripheral blood (PB) aliquots, (age range 1 to 97 years, median 60; 135 male, 98 female).

#### COVID-19 study samples

With the exceptions of age and gender investigators were blinded to all other demographics due to ethical constraints. Samples were less than 24 hours old and obtained from patients admitted to Leeds Teaching Hospitals NHS Trust between 7th April and 16th July 2020. Samples were collected from 52 patients: 1) (Mild), 20 samples from mild cases of COVID-19, defined by positive RT-PCR for SARS-CoV-2 and not requiring Intensive Care Unit (ITU) support (age range 20 to 90 years, median 73; 13 males, 7 females. 2) (**S**evere) 12 samples from severe cases of COVID-19, defined by positive RT-PCR for SARS-CoV-2 and requiring ITU support (age range 44 to 82 years, median 64; 6 males, 6 females). 3) (**C**ontrol) 20 control samples from patients with no features of COVID-19 and negative RT-PCR for SARS-CoV-2, 16/20 were not on ITU, 4/20 were on ITU (age range 20 to 91 years, median 57; 11 males, 9 females). Flow cytometry was performed, and plasma extracted using standard methods and stored at −20°C. CXCL16 ELISA was performed on 40 samples (12 C [including 2 ITU patients], 10 S, 18 M).

### Flow cytometry

Samples were analysed using 2 phenotyping panels (Supplementary Table 1). Consecutive sample analyses were performed using a FACS Canto II flow cytometer (BD Biosciences) verified through daily calibration with CS&T beads (BD Biosciences) and 8 peak Rainbow beads (Spherotech) utilising a 7-parameter panel (CD8-FITC, CD16-PE, CD4-PerCP-Cy5.5, CD3-PE-Cy7, CD8-APC, CD45-APC-Cy7, CD186-BV421 [Becton Dickinson]). Samples from COVID-19 patients and controls were assessed on a Cytoflex LX cytometer (Beckman Coulter), verified through daily calibration with CytoFLEX Daily QC Fluorospheres (Beckman Coulter), using a 12-parameter panel (CD57-FITC, CD4-PE, CD161-PC7, CD8-KrO, CD279-PC5.5, CD45RA-A700, CD3-APC-A750 [Beckman Coulter] and CD56-BV605, CD45-PerCP, CD186-BV421, CD27-BV786, HLA-DR-BUV395 [Becton Dickinson]). A minimum of 50,000 CD3^+^ T cell events were assessed and analysed using standard methods. All analyses were only performed once due to the volume of sample available. Data was analysed with Kaluza Analysis software version 2.1 (Beckman Coulter) and Cytobank software (Beckman Coulter) Representative gating strategy can be seen in Supplementary figure 1.

### ELISA

Plasma CXCL16 levels were measured in duplicate using a commercial ELISA (ThermoFisher Scientific), with a coefficient of variation of less than 10%, according to the manufacturer’s specifications.

### Statistics

Data was analysed with GraphPad Prism version 9.0.0 (GraphPad software). Categorical data was compared with Mann-Whitney statistical analyses. Spearman Rank and regression analysis was used to assess data correlation, p <0.05 was considered significant.

### Study approval

Local and National ethical approval (IRAS: 284369) allowed collection of anonymised excess peripheral blood from patients tested for SARS-CoV-2 infection.

## Author contributions

DP carried out the experiments, planned the study, analysed data, performed statistical analysis and wrote the manuscript. SD, RL acquired data. RP, SG, CM, GC, SR were involved in the design of the study. PH, TM, LA, KR, CM, SP were involved in the identification and collection of patient samples. RB and DN planned the study, interpreted the data and drafted the manuscript. All authors critically reviewed the manuscript.

## Acknowledgements

We would like to thank the HMDS department, University Hospitals of Leeds NHS Trust for partial funding of this project and use of facilities. Additionally, we want to thank Alison Bell, Markus Kaymer and Michael Kapinski from Beckman Coulter, Bev Goward from BD Biosciences and Thermofisher for support with reagents and technical assistance. We also like to thank Rachael Barlow, University of Leeds, for use of the ELISA plate reader, and Anne Gowing, Research and Innovation, University Hospitals of Leeds NHS Trust, who assisted with the ethics application process.

## References

1. Huang C, Wang Y, Li X, Ren L, Zhao J, Hu Y, et al. Clinical features of patients infected with 2019 novel coronavirus in Wuhan, China. Lancet. 2020;395(10223):497–506.

2. Wu Z, and McGoogan JM. Characteristics of and Important Lessons From the Coronavirus Disease 2019 (COVID-19) Outbreak in China: Summary of a Report of 72314 Cases From the Chinese Center for Disease Control and Prevention. JAMA. 2020;323(13):1239–42.

3. Vabret N, Britton GJ, Gruber C, Hegde S, Kim J, Kuksin M, et al. Immunology of COVID-19: Current State of the Science. Immunity. 2020;52(6):910–41.

4. Soy M, Keser G, Atagunduz P, Tabak F, Atagunduz I, and Kayhan S. Cytokine storm in COVID-19: pathogenesis and overview of anti-inflammatory agents used in treatment. Clin Rheumatol. 2020;39(7):2085–94.

5. Wiersinga WJ, Rhodes A, Cheng AC, Peacock SJ, and Prescott HC. Pathophysiology, Transmission, Diagnosis, and Treatment of Coronavirus Disease 2019 (COVID-19): A Review. JAMA. 2020;324(8):782–93.

6. Hanley B, Lucas SB, Youd E, Swift B, and Osborn M. Autopsy in suspected COVID-19 cases. J Clin Pathol. 2020;73(5):239–42.

7. Borczuk AC, Salvatore SP, Seshan SV, Patel SS, Bussel JB, Mostyka M, et al. COVID-19 pulmonary pathology: a multi-institutional autopsy cohort from Italy and New York City. Mod Pathol. 2020;33(11):2156–68.

8. Guaraldi G, Meschiari M, Cozzi-Lepri A, Milic J, Tonelli R, Menozzi M, et al. Tocilizumab in patients with severe COVID-19: a retrospective cohort study. Lancet Rheumatol. 2020;2(8):e474–e84.

9. Group RC, Horby P, Lim WS, Emberson JR, Mafham M, Bell JL, et al. Dexamethasone in Hospitalized Patients with Covid-19 - Preliminary Report. N Engl J Med. 2020.

10. Petrilli CM, Jones SA, Yang J, Rajagopalan H, O’Donnell L, Chernyak Y, et al. Factors associated with hospital admission and critical illness among 5279 people with coronavirus disease 2019 in New York City: prospective cohort study. BMJ. 2020;369:m1966.

11. Akbar AN, and Gilroy DW. Aging immunity may exacerbate COVID-19. Science. 2020;369(6501):256–7.

12. Lustig A, Carter A, Bertak D, Enika D, Vandanmagsar B, Wood W, et al. Transcriptome analysis of murine thymocytes reveals age-associated changes in thymic gene expression. Int J Med Sci. 2009;6(1):51–64.

13. Garner LC, Klenerman P, and Provine NM. Insights Into Mucosal-Associated Invariant T Cell Biology From Studies of Invariant Natural Killer T Cells. Front Immunol. 2018;9:1478.

14. Billerbeck E, Kang YH, Walker L, Lockstone H, Grafmueller S, Fleming V, et al. Analysis of CD161 expression on human CD8+ T cells defines a distinct functional subset with tissue-homing properties. Proc Natl Acad Sci U S A. 2010;107(7):3006–11.

15. Dusseaux M, Martin E, Serriari N, Peguillet I, Premel V, Louis D, et al. Human MAIT cells are xenobiotic-resistant, tissue-targeted, CD161hi IL-17-secreting T cells. Blood. 2011;117(4):1250–9.

16. Meftahi GH, Jangravi Z, Sahraei H, and Bahari Z. The possible pathophysiology mechanism of cytokine storm in elderly adults with COVID-19 infection: the contribution of “inflame-aging”. Inflamm Res. 2020;69(9):825–39.

17. Cunha LL, Perazzio SF, Azzi J, Cravedi P, and Riella LV. Remodeling of the Immune Response With Aging: Immunosenescence and Its Potential Impact on COVID-19 Immune Response. Front Immunol. 2020;11:1748.

18. Bonafe M, Prattichizzo F, Giuliani A, Storci G, Sabbatinelli J, and Olivieri F. Inflamm-aging: Why older men are the most susceptible to SARS-CoV-2 complicated outcomes. Cytokine Growth Factor Rev. 2020;53:33–7.

19. Yanez ND, Weiss NS, Romand JA, and Treggiari MM. COVID-19 mortality risk for older men and women. BMC Public Health. 2020;20(1):1742.

20. Diao B, Wang C, Tan Y, Chen X, Liu Y, Ning L, et al. Reduction and Functional Exhaustion of T Cells in Patients With Coronavirus Disease 2019 (COVID-19). Front Immunol. 2020;11:827.

21. Huang I, and Pranata R. Lymphopenia in severe coronavirus disease-2019 (COVID-19): systematic review and meta-analysis. J Intensive Care. 2020;8:36.

22. He Z, Zhao C, Dong Q, Zhuang H, Song S, Peng G, et al. Effects of severe acute respiratory syndrome (SARS) coronavirus infection on peripheral blood lymphocytes and their subsets. Int J Infect Dis. 2005;9(6):323–30.

23. Zhao Y, Zhao Z, Wang Y, Zhou Y, Ma Y, and Zuo W. Single-Cell RNA Expression Profiling of ACE2, the Receptor of SARS-CoV-2. Am J Respir Crit Care Med. 2020;202(5):756–9.

24. Day C, Patel R, Guillen C, and Wardlaw AJ. The Chemokine CXCL16 is Highly and Constitutively Expressed by Human Bronchial Epithelial Cells. Exp Lung Res. 2009;35(4):272–83.

25. Morgan AJ, Guillen C, Symon FA, Huynh TT, Berry MA, Entwisle JJ, et al. Expression of CXCR6 and its ligand CXCL16 in the lung in health and disease. Clin Exp Allergy. 2005;35(12):1572–80.

26. Ashhurst AS, Flórido M, Lin LCW, Quan D, Armitage E, Stifter SA, et al. CXCR6-Deficiency Improves the Control of Pulmonary Mycobacterium tuberculosis and Influenza Infection Independent of T-Lymphocyte Recruitment to the Lungs. Front Immunol. 2019;10.

27. Nakayama T, Hieshima K, Izawa D, Tatsumi Y, Kanamaru A, and Yoshie O. Cutting edge: profile of chemokine receptor expression on human plasma cells accounts for their efficient recruitment to target tissues. J Immunol. 2003;170(3):1136–40.

28. Wein AN, McMaster SR, Takamura S, Dunbar PR, Cartwright EK, Hayward SL, et al. CXCR6 regulates localization of tissue-resident memory CD8 T cells to the airways. J Exp Med. 2019;216(12):2748–62.

29. Tu GW, Ju MJ, Zheng YJ, Hao GW, Ma GG, Hou JY, et al. CXCL16/CXCR6 is involved in LPS-induced acute lung injury via P38 signalling. J Cell Mol Med. 2019;23(8):5380–9.

30. Izquierdo MC, Martin-Cleary C, Fernandez-Fernandez B, Elewa U, Sanchez-Nino MD, Carrero JJ, et al. CXCL16 in kidney and cardiovascular injury. Cytokine Growth Factor Rev. 2014;25(3):317–25.

31. Zhang W, Hua T, Li J, Zheng L, Wang Y, Xu M, et al. CXCL16 is activated by p-JNK and is involved in H2O2-induced HK-2 cell injury via p-ERK signaling. Am J Transl Res. 2018;10(11):3723–32.

32. Steffen S, Abraham S, Herbig M, Schmidt F, Blau K, Meisterfeld S, et al. Toll-Like Receptor-Mediated Upregulation of CXCL16 in Psoriasis Orchestrates Neutrophil Activation. J Invest Dermatol. 2018;138(2):344–54.

33. Gunther C, Carballido-Perrig N, Kaesler S, Carballido JM, and Biedermann T. CXCL16 and CXCR6 are upregulated in psoriasis and mediate cutaneous recruitment of human CD8+ T cells. J Invest Dermatol. 2012;132(3 Pt 1):626–34.

34. Chua RL, Lukassen S, Trump S, Hennig BP, Wendisch D, Pott F, et al. COVID-19 severity correlates with airway epithelium-immune cell interactions identified by single-cell analysis. Nat Biotechnol. 2020;38(8):970–9.

35. Picton ACP, Paximadis M, Chaisson RE, Martinson NA, and Tiemessen CT. CXCR6 gene characterization in two ethnically distinct South African populations and association with viraemic disease control in HIV-1-infected black South African individuals. Clin Immunol. 2017;180:69–79.

36. Patel P, Hiam L, Sowemimo A, Devakumar D, and McKee M. Ethnicity and covid-19. BMJ. 2020;369:m2282.

37. Parrot T, Gorin JB, Ponzetta A, Maleki KT, Kammann T, Emgard J, et al. MAIT cell activation and dynamics associated with COVID-19 disease severity. Sci Immunol. 2020;5(51).

38. Fergusson JR, Huhn MH, Swadling L, Walker LJ, Kurioka A, Llibre A, et al. CD161(int)CD8+ T cells: a novel population of highly functional, memory CD8+ T cells enriched within the gut. Mucosal Immunol. 2016;9(2):401–13.

39. Chiba A, Murayama G, and Miyake S. Mucosal-Associated Invariant T Cells in Autoimmune Diseases. Front Immunol. 2018;9:1333.

40. Gazali AM, Schroderus AM, Nanto-Salonen K, Rintamaki R, Pihlajamaki J, Knip M, et al. Mucosal-associated invariant T cell alterations during the development of human type 1 diabetes. Diabetologia. 2020;63(11):2396–409.

41. Liao M, Liu Y, Yuan J, Wen Y, Xu G, Zhao J, et al. Single-cell landscape of bronchoalveolar immune cells in patients with COVID-19. Nat Med. 2020;26(6):842–4.

42. Yu H, Yang A, Liu L, Mak JYW, Fairlie DP, and Cowley S. CXCL16 Stimulates Antigen-Induced MAIT Cell Accumulation but Trafficking During Lung Infection Is CXCR6-Independent. Front Immunol. 2020;11:1773.

43. Sandor AM, Jacobelli J, and Friedman RS. Immune cell trafficking to the islets during type 1 diabetes. Clin Exp Immunol. 2019;198(3):314–25.

44. Xiao X, and Cai J.Mucosal-Associated Invariant T Cells: New Insights into Antigen Recognition and Activation. Front Immunol. 2017;8:1540.

45. Kang CK, Han GC, Kim M, Kim G, Shin HM, Song KH, et al. Aberrant hyperactivation of cytotoxic T-cell as a potential determinant of COVID-19 severity. Int J Infect Dis. 2020;97:313–21.

46. Jiang Y, Wei X, Guan J, Qin S, Wang Z, Lu H, et al. COVID-19 pneumonia: CD8(+) T and NK cells are decreased in number but compensatory increased in cytotoxic potential. Clin Immunol. 2020;218:108516.

47. Lu B, Liu M, Wang J, Fan H, Yang D, Zhang L, et al. IL-17 production by tissueresident MAIT cells is locally induced in children with pneumonia. Mucosal Immunol. 2020;13(5):824–35.

48. De Biasi S, Meschiari M, Gibellini L, Bellinazzi C, Borella R, Fidanza L, et al. Marked T cell activation, senescence, exhaustion and skewing towards TH17 in patients with COVID-19 pneumonia. Nat Commun. 2020;11(1):3434.

49. Kuri-Cervantes L, Pampena MB, Meng W, Rosenfeld AM, Ittner CAG, Weisman AR, et al. Comprehensive mapping of immune perturbations associated with severe COVID-19. Sci Immunol. 2020;5(49).

